# Ventral Pallidum is Essential for Cocaine Reinstatement After Voluntary Abstinence

**DOI:** 10.1101/653741

**Authors:** Mitchell R. Farrell, Christina M. Ruiz, Erik Castillo, Lauren Faget, Christine Khanbijian, Siyu Liu, Hannah Schoch, Gerardo Rojas, Thomas S. Hnasko, Stephen V. Mahler

## Abstract

Addiction is a chronic relapsing disorder, and during recovery many people experience several relapse events as they attempt to voluntarily abstain from drug. New preclinical relapse models have emerged which capture this common human experience of relapse after voluntary abstinence, and mounting evidence indicates that reinstatement of drug seeking after voluntary abstinence recruits neural circuits distinct from reinstatement following experimenter-imposed abstinence, or abstinence due to extinction training. Ventral pallidum (VP), a key limbic node involved in drug seeking, has well-established roles in conventional reinstatement models tested following extinction training, but it is unclear whether this region also participates in more translationally-relevant models of relapse. Here we show that chemogenetic inhibition of VP neurons strongly attenuates cocaine-, context-, and cue-induced reinstatement tested after voluntary, punishment-induced abstinence. This effect was strongest in the most compulsive, punishment-resistant rats, and reinstatement was associated with neural activity in anatomically-defined VP subregions. VP inhibition also attenuated the propensity of rats to display ‘hesitations,’ a risk assessment behavior seen during punished drug taking that is likely due to concurrent approach and avoidance motivations. These results indicate that VP, unlike other connected limbic brain regions, is essential for reinstatement of drug seeking after voluntary abstinence. Since VP inhibition effects were strongest in the most compulsively cocaine-seeking individuals, this could indicate that VP plays a particularly important role in the most pathological, addiction-like behavior, making it an attractive target for future therapeutic interventions.

## Introduction

Addiction is characterized by persistent drug use despite negative consequences, and a lasting vulnerability to relapse after protracted periods of abstinence [1–3]. Typically, human addicts eventually recognize the negative consequences of their behavior, and choose to cease using drugs—a decision they usually renege upon when tempted by drug cues, small doses of drug, or stressors [4]. In rodent relapse models, reinstatement of seeking is triggered by analogous stimuli, usually following a period of imposed abstinence from drug (incubation), or explicit extinction training. Recently, voluntary abstinence-based rodent models have emerged, capturing the fact that in many cases addicted people choose to stop using drugs due to mounting negative life consequences, rather than due to extinction training or external forces [5–9]. This is important because in rodents, the neural substrates underlying reinstatement differ based upon how abstinence was achieved, be it experimenter-imposed, through extinction training, or through voluntary cessation due to punishment or availability of more attractive alternative reinforcers [10–15]. If the brain substrates of human relapse similarly depend upon why a person stopped using drugs, then considering these factors in preclinical models will be essential for developing effective interventions to treat addiction.

A hallmark of addiction is an inability to limit drug intake in the face of negative life consequences. This can be modeled in rodents by training them to self-administer drugs, then introducing consequences to continued use, such as co-delivered footshock [5, 6, 16–20]. As in humans, most rodents readily suppress their drug intake when negative outcomes begin to result from their drug use. However, a subset of rodents show punishment-resistant drug intake [17, 21–23], similar to the proportion of humans who use drugs that ultimately become addicted [24]. Punishment-resistant animals also exhibit the most robust reinstatement of cocaine and methamphetamine seeking [17, 25], suggesting that compulsive use and liability to relapse involve common underlying neural mechanisms. Indeed, the circuitry underlying compulsive cocaine intake overlaps with the limbic substrates of reinstatement behavior, at least when tested following extinction training [23, 26–29].

One brain region that has emerged as being crucial for motivated behavior is the ventral pallidum (VP), the main efferent target of nucleus accumbens [30–35]. VP is thought to help translate motivation into action [36–39], and accordingly, VP neural activity encodes reward motivation in rodents, monkeys, and humans [40–42], including for cocaine [43]. VP is also required for seeking of several abused drugs [44–49], and for cocaine reinstatement triggered by cues, stress, or cocaine following extinction training [47, 50, 51]. Notably, VP is a heterogeneous structure, with functionally and anatomically distinct dorsolateral/ventromedial, and rostral/caudal subregions that mediate distinct aspects of reward seeking, including cocaine reinstatement [43, 47, 52–59]. Given these results, and recent findings that VP contains phenotypically distinct populations of reward- and aversion-related neurons [30–32, 60–62], its role in drug seeking under translationally-relevant mixed motivation circumstances was of interest to us.

Here we explore effects of transiently and reversibly inhibiting VP neurons of punishment-resistant or punishment-sensitive rats with designer receptors (DREADDs) [63], determining effects on punished cocaine seeking, context, discrete cue, and primed reinstatement after voluntary abstinence, as well as on cocaine-induced locomotion. We also assessed reinstatement-related Fos in VP subregions. These studies shed light on the functions of this essential, but understudied nucleus within cocaine addiction-related neural circuits.

## Methods

### Subjects

Male (*n*=50) and female (*n*=36) Long-Evans rats (220-250g at the start of experiments) were bred at the University of California Irvine or obtained from Envigo, and were pair housed on a 12hr reverse light/dark cycle with *ad libitum* food and water for all experiments. All training and testing was conducted in the dark period. Procedures were approved by the UCI Institutional Animal Care and Use Committee, and are in accordance with the NIH Guide for the Care and Use of Laboratory Animals [64].

### Surgery

Animals were anesthetized with ketamine (56.5mg/kg), xylazine (8.7mg/kg), and the non-opioid analgesic meloxicam (1.0mg/kg), and implanted with indwelling jugular catheters exiting the dorsal back. In the same surgery, they also received bilateral viral vector injections (250-300nL) into VP with pressure injections using a Picospritzer and glass micropipette. See Figure 1 for schematic of procedures.

**Figure 1.**
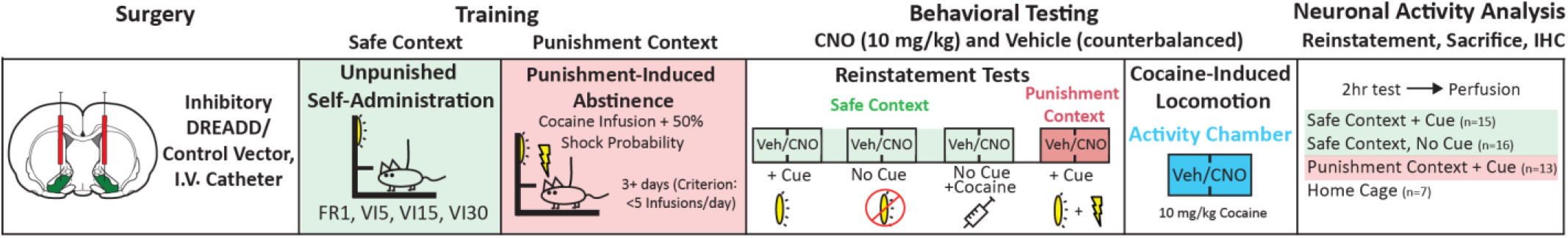
Schematic of experimental timeline. Following DREADD or control AAV injection, rats underwent cocaine self-administration, punishment training, followed by reinstatement and cocaine-induce locomotor testing. A final reinstatement test preceded sacrifice for neuronal activity (Fos) analysis.

### Viral Constructs

To transduce VP neurons with hM4Di inhibitory DREADDs, we used a human synapsin (hSyn) promoter-driven AAV with mCitrine (*n*=44; U North Carolina vector core: *AAV2-hSyn-HA-hM4D(Gi)-IRES-mCitrine)* or mCherry (*n*=16; Addgene: *AAV2-hSyn-hM4D(Gi)-mCherry*) reporter. To control for non-specific impact of viral transduction and clozapine-N-oxide (CNO) in the absence of DREADDs, an eGFP-only reporter without DREADDs (*n*=7; Addgene: *AAV2-hSyn-eGFP)*, was employed in a group of control rats [65–67].

### Anatomical Analysis of DREADD Expression

hM4Di DREADD/reporter expression was visualized with immunofluorescent amplification, with Substance P co-stain allowing demarcation of VP borders (see Table S1 for detailed list of antibodies). Rats with at least 40% of VP volume expressing DREADDs/reporter, and at least 40% of virus expression localized within VP borders were considered hits (*n*=46). Rats with more than 60% of DREADD expression localized outside VP (quantified blind to group and behavioral results) were considered misses (*n*=13). Since rats with extra-VP DREADD expression did not behaviorally differ from fluorophore-only rats on reinstatement tests (no main effect of group or CNO treatment on reinstatement types, *Fs*<1.29, *ps*>0.27; Fig. S1), they were combined into a single control group (*n*=20) for subsequent analyses of CNO effects in the absence of VP DREADDs.

### RNAscope Analysis of DREADD Expression in VP Neurons

PFA-fixed brains were serially cut (16μm) on a cryostat and mounted directly onto glass slides. Sections were stored at –80°C until processing for RNAscope Multiplex Fluorescent assay (Advanced Cell Diagnostics). Briefly, sections were warmed on a hot plate for 30 minutes at 60°C then boiled at 100° for 6 min in target retrieval solution. Sections were then dehydrated in 100% ethanol and treated with protease (pretreatment reagents, cat. No. 322380). RNA hybridization probes included antisense probes against rat *gad1* (316401-C1) and *slc17a6* (317011-C3), respectively labeled with alexa488 and atto647 fluorophores. Slides were then incubated with rabbit anti-DsRed primary antibodies (632496, Clontech) and donkey anti-rabbit AlexaFluor 594 secondary antibodies (711-585-152, Jackson ImmunoResearch), counterstained with DAPI, and coverslipped using Fluoromount-G mounting medium. Images for cell counting were taken at 63x (1.4 NA) magnification using a Zeiss AxioObserver Z1 widefield Epifluorescence microscope with a Zeiss ApoTome 2.0 for structured illumination and Zen Blue software. An average of 186 +/− 11 cells positive for AAV-hSyn-hM4D(Gi)-mCherry were counted per brain (*n*=3 rats).

### Drugs

Cocaine HCl (NIDA) was dissolved in 0.9% saline. Cocaine was available for self-administration at 0.2mg/50μL infusion for male rats, and 0.15mg/50μL infusion for female rats [68, 69]. Cocaine (10mg/kg) was used for primed reinstatement and locomotion testing. CNO was dissolved in a vehicle of 5% DMSO in 0.9% saline, and injected i.p. at 10mg/kg 30min prior to behavioral testing.

### Behavioral Testing Apparatus

Self-administration training and testing took place in Med Associates operant chambers within sound-attenuating boxes, equipped with two retractable levers with white lights above them, and a tone generator. Cocaine-induced locomotion testing was conducted in 43×43×30.5cm Med Associates locomotor testing chambers.

### Self-Administration Training in Safe Context

We employed a punishment-induced abstinence/reinstatement protocol modeled after previous reports [5, 20]. Initial self-administration occurred in a ‘safe context,’ signaled by presence of a white or red house light, peppermint or orange scent, and plain or polka dot pattern walls (randomly assigned). Intravenous cocaine was administered via a pump located outside the sound-attenuating box. Rats received five daily 2hr sessions of fixed-ratio 1 (FR1) training where an active lever press delivered a 3.6sec cocaine infusion, and concurrent stimulus light + 2.9-kHz tone. A 20sec timeout period (signaled by dimming of the house light) followed each infusion/cue presentation, during which additional lever presses did not yield cocaine delivery. Pressing on an inactive lever was recorded but had no consequences. Following FR1 training to criterion (>10 infusions), rats then completed 3 days of variable-interval 5 schedule (VI5), on which an active lever press initiated a timer with an average duration of 5sec, and another press after that interval delivered a cocaine infusion with the light+tone cue. The VI schedule was increased to VI15 for the next 3 days, then VI30 for an additional 3-6 days until rats had stable performance (Fig. S2).

### Punishment Context Testing and Training

Following safe context self-administration training, rats (*n*=35) began punishment training in a distinct chamber, with the opposite context used for safe context training. Rats continued on a VI30 schedule, but 50% of cocaine infusions/cues were accompanied by a 0.3mA foot shock (0.5sec). To test effects of inhibiting VP during punished cocaine intake, a subset of animals were injected with CNO (*n*=22) or vehicle (*n*=13) prior to each of two daily shock punishment (0.3 mA) training sessions. In a crossover design, these rats were administered the opposite treatment (vehicle/CNO) prior to a third punished intake session 48hrs later, then a fourth punished cocaine intake training session with no vehicle or CNO injection 24hrs later. Another group of rats (*n*=31) received no injections during punishment context training. After 3-4 days of shock training at 0.30mA, shock increased by 0.15mA every 2 training days, up to 0.75mA, until voluntary abstinence criterion was met in all rats (<5 active lever presses for 2 consecutive days). Sensitivity to punishment was determined in two ways. A suppression ratio (infusions on day 1 punishment/infusions on last day unpunished; [17, 70]) was calculated as a measure of initial punishment sensitivity, with high ratios reflecting relative insensitivity to shock-suppression of intake. Rats were also coded based on the maximum level of shock they tolerated during punishment training, before meeting abstinence criterion.

### Measuring Mixed Motivations During Punished Cocaine Intake: “Hesitation” Behavior

During punished cocaine intake training sessions, rats exhibited a species-typical risk assessment behavior [71, 72] we term ‘hesitation behavior,’ in which they stretch their trunk and extend their forepaw towards the active or inactive lever, but rapidly retract it without completing the press to deliver cocaine and probabilistic shock. Hesitations directed at the active and inactive levers were quantified using video analysis by a blinded observer on the final day of safe context self-administration, and the first day of punishment context self-administration in CNO- and vehicle-treated rats.

### Reinstatement Tests

A series of 2hr reinstatement tests commenced 48hrs after rats met abstinence criterion, with 48hrs elapsing between each test. Reinstatement tests occurred in: the safe context with response contingent cues (*n*=66; vehicle and CNO administered on separate days in counterbalanced order), the safe context without cues (vehicle/CNO, *n*=31), the safe context with no cues immediately after a cocaine priming injection (10mg/kg; vehicle/CNO, *n*=38), the punishment context with cues (vehicle/CNO, *n*=35), and the punishment context without cues (vehicle only, *n*=24). For reinstatement tests with cues, active lever presses yielded 3.6sec cocaine-associated cue presentations (with no cocaine or shock), delivered on a VI-30 schedule, and followed by a 15sec timeout period. For tests without discrete cues, lever presses were inconsequential, but were recorded.

### Cocaine Induced-Locomotion

Following reinstatement tests, a subset of rats (*n*=51) were habituated to a locomotor testing chamber for 2 consecutive days, followed by two 2hr locomotor tests, 48hrs apart. Next, rats were immediately placed in the chamber for 30min after vehicle/CNO, injected with cocaine, and returned to the chamber for 90min. Horizontal locomotor activity and vertical rearing were recorded via infrared beam breaks.

### Reinstatement-Related Fos

To examine VP neuronal activity during reinstatement, rats underwent a final drug-free 2hr reinstatement test, 48hr after their last vehicle/CNO reinstatement test. They were tested in one of the following reinstatement types: the safe context with response contingent cues (*n*=15), the safe context without cues (*n*=16), the punishment context with cues (*n*=13), or no reinstatement (removed directly from their home cage after equivalent self-administration/reinstatement training, *n*=7). After the 2hr test, rats were returned to their home cages for 1hr, then perfused with saline (0.9%) and paraformaldehyde (4%), and brains sectioned (40μm) following cryoprotection in 20% sucrose azide.

### Fos Quantification

To allow Fos quantification within anatomically-defined VP subregions, we stained for Fos + substance P to define VP borders. Ventromedial, ventrolateral, and dorsolateral subregions of substance P-defined VP were delineated with reference to adjacent sections stained for substance P and neurotensin, defining ventromedial/dorsolateral VP [54, 73] (Table S1). Images of VP were taken at 5x magnification, and one section/animal was quantified bilaterally in rostral VP (+0.12 to +0.60 mm relative to Bregma), and another in caudal VP (−0.48 to −0.24 mm; [74]). Fos+ neurons were identified using the Stereoinvestigator (Microbrightfield) particle counter tool with thresholding parameters incorporating particle size (average size 100μm^2^), minimum distance between nuclei (150μm), and color relative to background. Fos density (Fos/mm^2^) was computed for each VP subregion on each slice (average of both hemispheres) of each rat. All structure delineation and quantification was done blind to experimental conditions, and imaging/analysis settings were consistent across animals.

### Statistical Analyses

Effects of punishment on self-administration were examined with repeated measures ANOVAs, including day (1 unpunished, 3 punished days) and behavioral output (active lever, inactive lever, infusions) factors. Punishment sensitive versus resistant groups were compared on reinstatement with a two-way ANOVA with punishment sensitivity group and reinstatement type as factors. Pearson correlation was used for assessing relationships between suppression ratio and reinstatement behavior. Effects of punishment sensitivity group on unpunished cocaine intake and cocaine-induced locomotion were examined with unpaired *t*-tests. Effects of CNO in control and VP-hM4Di rats on hesitation behavior were computed with one-way ANOVA. Effects of CNO on each reinstatement type in VP hM4Di-expressing and control rats were examined using separate repeated measures ANOVAs with drug (vehicle/CNO) and lever (active/inactive) factors. Effects of VP inhibition on reinstatement in punishment resistant and punishment sensitive rats were computed as change from vehicle day behavior (CNO-vehicle), and compared with unpaired *t*-test. Separate one-way ANOVAs compared behavioral groups on Fos in each VP subregion. Effects of rostrocaudal VP location on Fos was examined with a two-way ANOVA with rostral/caudal site, and reinstatement type factors. Separate two-way ANOVAs were used to compare CNO effects on cocaine-induced horizontal distance and rearing in control and VP-hM4Di rats. Tukey and Bonferroni corrected *t*-tests were used for posthoc comparisons as appropriate.

## Results

### Unpunished Self-Administration

Rats readily discriminated between the inactive and active lever (Lever: *F*_(1, 130)_=55.3, *p*<0.0001), and daily cocaine intake was stable by the final 3 days of training (*F*_(2, 130)_=0.87, *p*=0.42; Fig. S2). Male and female rats did not differ in active lever presses or sex-adjusted cocaine doses self-administered during the last 3d of training (no main effect of sex (lever: *F*_(1, 64)_=1.8, *p*=0.19, infusions: *F*_(1, 64)_=0.29, *p*=0.59) or day X sex interaction (Lever: *F* _(2, 128)_=1.0, *p*=0.37, infusion: *F*_(2, 128)_=0.48, *p*=0.62).

### Individual Differences in Cocaine Seeking under Punishment

As expected, cocaine-coincident shock (50% of infusions) in the punishment context suppressed cocaine self-administration overall (Day: *F*_(3, 585)_=30.1, *p*<0.0001, Fig. 2A). Most rats (80.3%; *n*=53) reached suppression criterion at the two lowest shock intensities (0.30-0.45mA: ‘punishment sensitive’ rats), but a subset of rats (19.7%, *n*=13) persisted in responding up to higher shock intensities (0.60-0.75mA: ‘punishment resistant’ rats; Fig. 2B-C). In addition, punishment resistant rats had higher suppression ratios (infusions on the first day in the punishment context/infusions on the last day in the safe context [17, 70]; mean ± SEM = 0.48 ± 0.09) than punishment sensitive rats (mean ± SEM = 0.25 ± 0.02; *t*_64_=4.4, p<0.0001). Notably, of the 13 punishment resistant rats in this study, 8 were female (26.7% of tested females), while 5 were male (13.9% of tested males).

**Figure 2.**
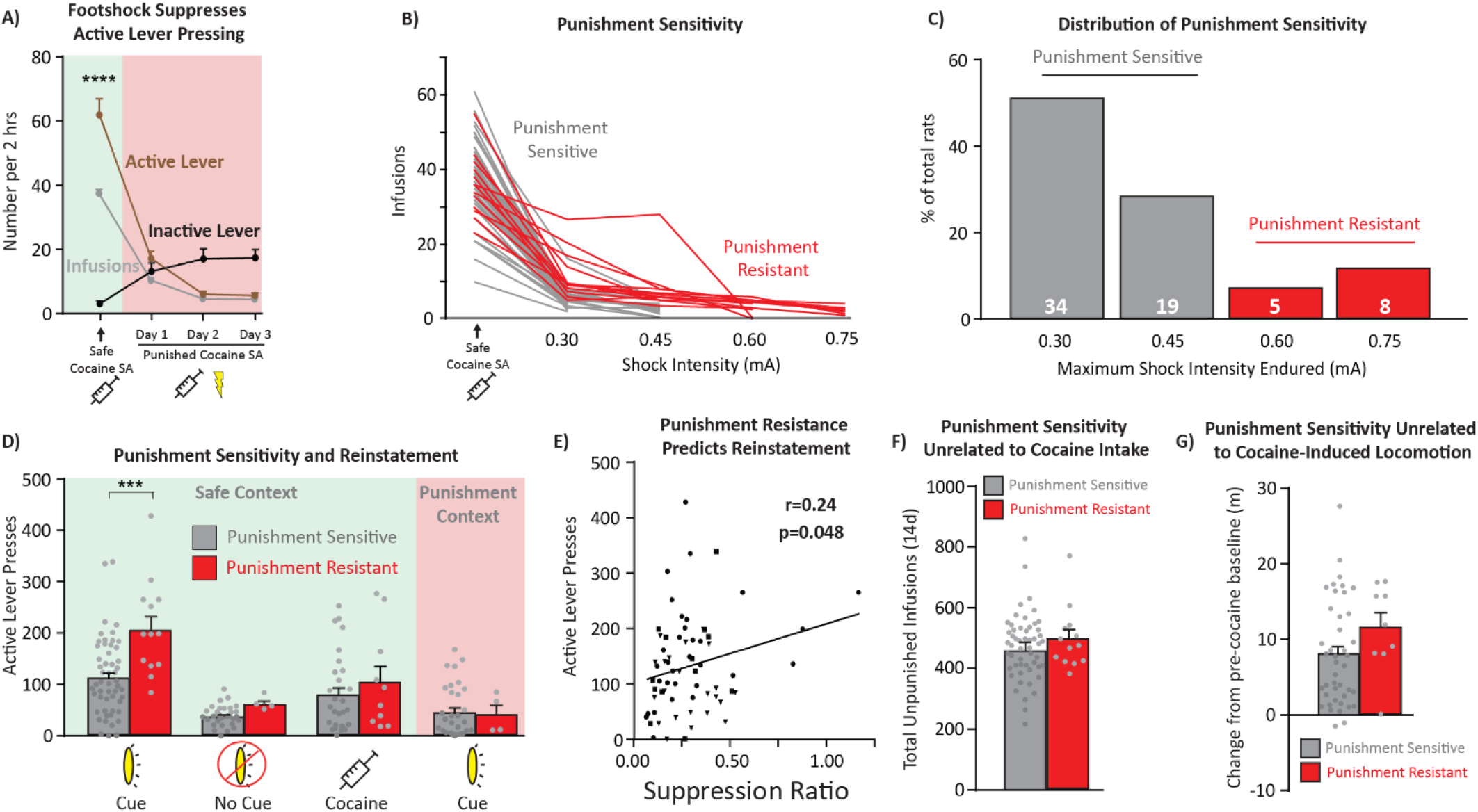
Punishment-resistant rats are more prone than punishment sensitive rats to cue, but not cocaine primed reinstatement. **A)** Probabilistic footshock reduces active lever pressing across all rats, while increasing the number of inactive presses. SA = self-administration. **B)** Individual variation in punishment sensitivity. Rats that reached at least 0.60mA footshock were considered punishment resistant (red). Punishment sensitive rats (gray) stopped taking cocaine at <0.60mA footshock intensities. **C)** Distribution of punishment resistance. Most rats cease cocaine intake at low shock levels (Punishment Sensitive: gray): 53/66 rats, 80.3%), but a subset reached the highest shock levels (Punishment Resistant: red): 13/66 rats, 19.7%). **D)** Punishment resistant rats reinstated more in the safe context with cues relative to punishment sensitive rats. This effect was specific to this reinstatement condition, and punishment sensitivity did not relate to context-only or cocaine primed reinstatment. **E)** Punishment effects on cocaine taking as quantified with suppression ratio [17,70] correlated with the degree of reinstatement in the safe context with cues. **F-G)** Punishment resistant rats were no different than sensitive rats on cocaine intake (F) or cocaine-induced locomotion (G). *** p<0.001. Panel E, squares = vehicle injection, triangles = CNO injection, circles = no injection.

### Punishment Resistant Rats Reinstated More

Punishment resistant rats, once they received shock intensities high enough to suppress even their seeking, showed greater cue-induced reinstatement than punishment sensitive rats. However, this was only true in the “safe,” unpunished context, and not when response contingent cues were delivered in the “punishment” context (punishment sensitivity x reinstatement type interaction: *F*_(3, 166)_=2.91, *p*=0.036; punishment resistant vs. sensitive in safe context with cues: *t*_166_=4.63, *p*<0.0001; Fig. 2D). Punishment suppression ratio also correlated with the magnitude of cue reinstatement in the safe context (*r*=0.24, *p*=0.048; Fig. 2E), further supporting a relationship between shock resistance and reinstatement propensity.

Punishment resistance was unrelated to total prior cocaine infusions (punishment resistant vs. sensitive total unpunished infusions: *t*_64_=0.67, *p*=0.50; Fig. 2F), or to cocaine’s locomotor stimulating or reinstating effects (horizontal distance traveled: *t*_49_=1.45, *p*=0.15; Fig. 2G; rearing: *t*_49_=1.77, *p*=0.084; cocaine primed reinstatement: *t*_36_=0.83, *p*=0.41), indicating that punishment resistance and cue-induced reinstatement likely involve underlying individual differences in addiction-like compulsive cocaine seeking, rather than sensitivity to cocaine’s effects per se.

### DREADD Expression in VP Neuronal Populations

Robust hM4Di-DREADD expression was observed throughout the rostrocaudal extent of VP in this study (Fig. 3A-B). Fluorescent *in situ* hybridization (RNAscope) revealed colocalization of hM4Di expression with *gad1*^+^ neurons (85.3 +/− 3.2 %), with a smaller percentage colocalizing with *gad1*^−^ neurons (14.7 +/− 3.2 %; Fig. 3C-E), consistent with unbiased transfection of all VP neurons, as GABA neurons represent the predominant neuronal phenotype in VP [31]. Non-GABA neurons likely represent intermingled glutamatergic and cholinergic cell-types [31, 54].

**Figure 3.**
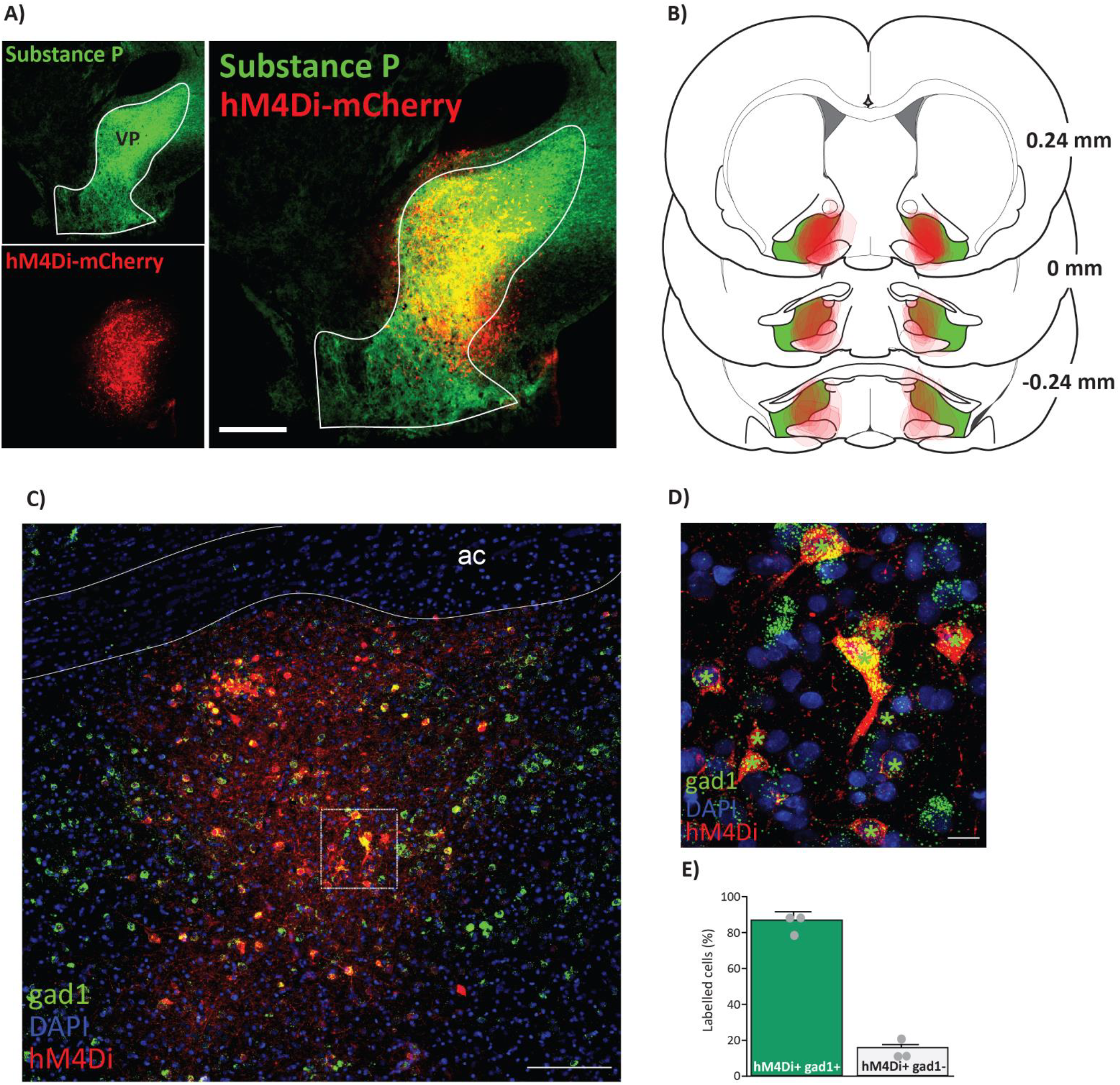
Inhibitory DREADD localization in VP. **A)** Immunofluorescent co-stain for hM4Di-mCherry (red) within Substance P-expressing VP borders (green). **B)** Virus expression for individual animals on rostrocaudal VP axis. **C**) Wide-field and **D)** high magnification images of fluorescent *in situ* hybridization for *gad1* (green) combined with immunofluorescence for hM4Di-mCherry (red). Co-positive hM4Di^+^ *gad1*^+^ cells labeled with green stars. **E)** 85.3 +/− 3.2 % of hM4Di^+^ cells are co-positive for *gad1.* 14.7 +/− 3.2 % of hM4Di^+^ cells are *gad1*^−^ (n=3 rats, total of 559 hM4Di^+^ cells counted). VM = ventromedial, VL = ventrolateral, DL = dorsolateral. AC = anterior commissure. Scale bars = 500μm (B), 200μm (C), and 20μm (D).

### CNO Effects on Punishment-Induced Suppression of Cocaine Intake in VP-hM4Di rats

On day 1 of punished cocaine self-administration, CNO in VP-hM4Di rats modestly, but non-significantly, decreased the number of active and inactive lever presses relative to control rats (Treatment: *F*_(1, 33)_=3.41, *p*=0.073; Fig. 4A), with no interaction of Treatment x Lever (*F*_(1, 33)_=0.27, *p*=0.61). CNO had no effect on lever pressing on day 2, though rats decreased their responding relative to day 1 (Day: *F*_(1, 33)_=20.56, *p*<0.0001). When vehicle and CNO treatments were reversed on punishment day 3, no further changes were observed (*p*>0.05).

**Figure 4.**
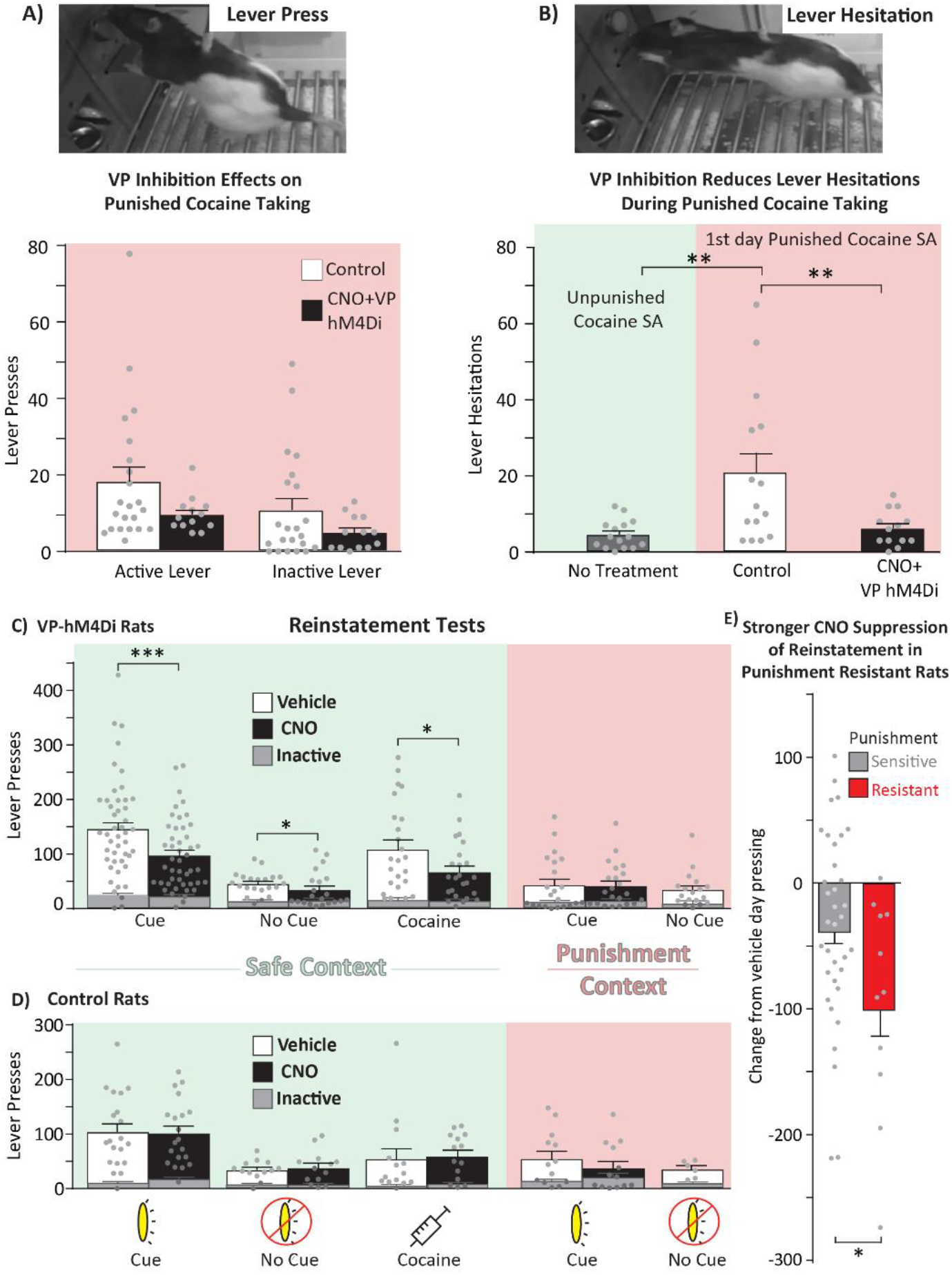
VP inhibition reduces reinstatement especially in punishment resistant rats. **A)** Top panel: Example picture of a standard lever press. Bottom panel: CNO in VP-hM4Di rats modestly reduces active and inactive lever pressing for cocaine under threat of punishment. Control = vehicle-injected rats and CNO-injected misses. **B)** Top panel: Hesitation behavior in which the rat stretches its trunk towards the lever and extends its paw without depressing the lever. Bottom panel: Active and inactive lever hesitations quantified during safe (green shading) and punished (red shading). CNO in VP-hM4Di rats reduced hesitations to relative to control rats. VP inhibition reduced hesitations to unpunished levels. Control = vehicle-injected rats and CNO-injected misses. **C)** Within subjects comparisons of reinstatement for VP-hM4Di rats in safe (green shading) and punishment (red shading) contexts. CNO in VP-hM4Di rats reduced reinstatement in the safe context with cues, without cues, and with cocaine and no cues, but not in the punishment context with cues. **D)** CNO in control rats did not affect reinstatement under any condition. Control = eGFP-only rats and rats with hM4Di expression primarily outside VP. **E)** CNO in VP-hM4Di punishment resistant rats (red bars) elicited a greater decrease in reinstatement relative to punishment sensitive rats (gray bars). Data presented as change from vehicle test baseline. *p*<0.05*, *p*<0.01**, *p*<0.001***.

Footshock+cocaine also increased “hesitation” behaviors targeted toward both the active and inactive levers, relative to the low levels seen during unpunished self-administration in the same context (*F*_(2, 40)_=7.93, *p*=0.0013; self-administration vs. vehicle: *p*=0.0022; Fig. 4B). Hesitations were highest in rats with the most active lever presses (correlation of hesitations and presses after both CNO; *r*=0.61, *p*=0.027; and vehicle; *r*=0.61, *p*=0.016), suggesting that hesitations may be a sensitive measure of deliberation about pursuing the now dangerous cocaine. Accordingly, CNO strongly suppressed hesitations, returning them to unpunished levels in VP-hM4Di rats (*p*=0.0013; CNO vs. vehicle: *p*=0.0082; Fig. 4B).

### VP DREADD Inhibition Suppresses Cocaine Reinstatement After Voluntary Abstinence

CNO in VP-hM4Di rats robustly suppressed context only-, cue-, and cocaine-induced reinstatement in the safe context, but failed to do so in control rats without VP DREADDs. In VP-hM4Di rats, CNO (compared to vehicle) reduced cue-induced active, but not inactive lever pressing in the safe context (Drug x Lever interaction: *F*_(1, 45)_=18.53, *p*<0.0001; Fig. 4C), and also suppressed safe context pressing without response-contingent cues (Drug x Lever interaction: *F*_(1, 20)_=4.31, *p*=0.05; Fig. 4C). Similarly, cocaine primed reinstatement (no cues) in the safe context was also suppressed by CNO in VP hM4Di rats (Drug x Lever interaction: *F*_(1, 23)_=7.94, *p*=0.01; Fig. 4C). Although we previously showed that rostral and caudal VP differentially mediate cue- and primed reinstatement [47], in these experiments our viral infection spanned most of the rostrocaudal axis of VP. In contrast to the safe context, CNO in VP-hM4Di rats failed to reduce cue-induced reinstatement in the punishment context (Drug x Lever interaction: *F*_(1, 21)_=0.19, *p*=0.66; Fig. 4C). In control rats without VP DREADDs, CNO had no effects on lever pressing in any reinstatement test (*p* values>0.05; Fig. 4D), suggesting that CNO effects here, as previously shown [45, 47, 75, 76], were specific to VP DREADD-expressing rats.

### VP Inhibition Suppressed Reinstatement the Most in Punishment-Resistant Rats

VP inhibition reduced safe context cue-induced reinstatement more in punishment-resistant rats than punishment-sensitive rats (*t*_44_=2.23, *p*=0.031; Fig. 4E). This effect was specific to the safe context with cues, as there was no such effect on other reinstatement types (*t* scores<1.26, *ps*>0.22). This finding suggests that VP plays an especially important role in relapse after punishment-imposed abstinence for the individual rats showing the most addiction-like behavior.

### VP Inhibition did not Affect Cocaine-Induced Locomotion

CNO failed to affect the locomotor-activating effects of cocaine in either VP-hM4Di or control groups (Treatment: *F*_(1, 49)_=0.63, *p*=0.43; treatment x group interaction: *F*_(1, 49)_=0.58, *p*=0.45; Fig. S3A), though it did reduce rearing behavior after cocaine in VP-hM4Di rats, but not controls (treatment x group interaction: *F*_(1, 49)_=10.24, *p*=0.0024; Fig. S3B), further suggesting specificity of these findings to cocaine seeking in particular. Moreover, CNO did not differentially reduce horizontal locomotion or rearing behavior in punishment-sensitive versus punishment-resistant VP-hM4Di rats (group x treatment interaction; locomotion: *F*_(3, 93)_=0.70, *p*=0.55; Rearing: *F*_(3, 93)_=0.61, *p*=0.61).

### VP Subregion Fos Recruited During Reinstatement

Relative to cocaine/shock-experienced rats sacrificed from their homecages, VP subregions showed strong Fos activation during all tested reinstatement conditions (*F*_(3, 47)_=3.93, *p*=0.014; punishment+cues, *p*=0.013; safe+cues, *p*=0.019; safe+no cues, *p*=0.043). Ventromedial VP was selectively activated (relative to home cage) by the punishment context with cues, but not by either safe context reinstatement test (*F*_(3, 47)_=2.67, *p*=0.05; punishment+cues: *p*=0.048; safe+cues: *p*=0.28; safe+no cues: *p*=0.09; Fig. 5A-C). In contrast, ventrolateral and dorsolateral VP were activated in all reinstatement conditions relative to homecage controls (ventrolateral: *F*_(3, 47)_=5.98, *p*=0.0015; safe+cues: *p*=0.0051; safe+no cues: *p*=0.0011; punishment+cues: *p*=0.0049; Fig. 5D; dorsolateral: *F*_(3, 47)_=4.63, *p*=0.006; home cage vs. safe+cues: *p*=0.0043; safe+no cues: *p*=0.017; punishment+cues: *p*=0.021; Fig. 5E). We then examined rostral and caudal sections of VP, given known rostrocaudal functional and anatomical differences [47, 54–56]. Overall, rostral VP had greater Fos density than caudal VP (*F*_(3, 47)_=4.8, *p*=0.0051), though this did significantly differ between reinstatement types (no reinstatement type X rostrocaudal position interaction; *F*_(3, 47)_=1.42, *p*=0.25; Fig. 5F).

**Figure 5.**
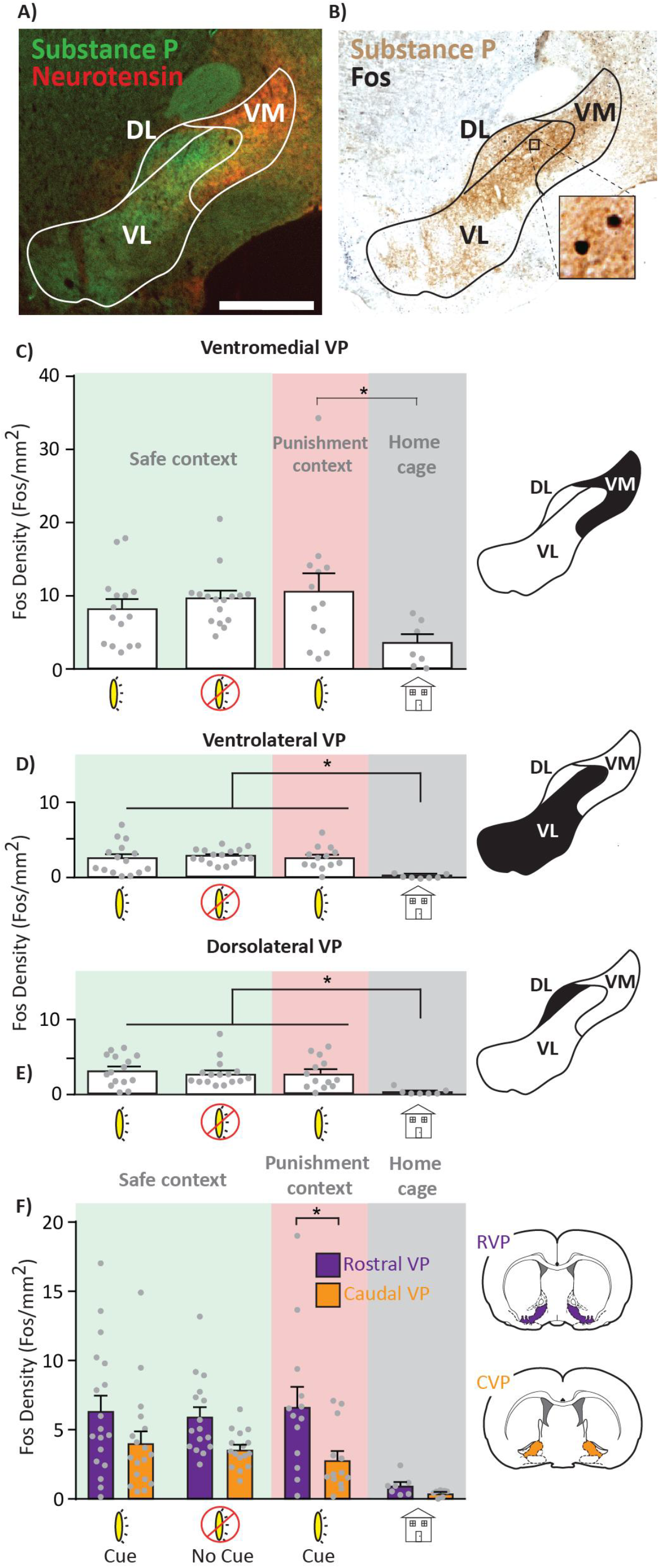
VP subregion Fos expression after reinstatement. **A)** Representative image of substance P (VP borders) and neurotensin (ventromedial VP marker) immunofluorescent co-stain. **B)** Representative image of substance P and Fos co-stained for Fos quantification. **C-E)** Comparison of ventromedial **(C)**, ventrolateral **(D)**, and dorsolateral **(E)** Fos density across reinstatement conditions. Green shading = safe context, red shading = punishment context, gray shading = home cage controls. **F)** Rostrocaudal Fos density across reinstatement conditions. Rostral VP reached significant only in the punishment context with cues. *p*<0.05*; VM = ventromedial, VL = ventrolateral, DL = dorsolateral. AC = anterior commissure. Scale bar = 1000μm.

### Sex Differences

Few sex differences with regard to VP manipulations were detected. Male and female rats exhibited comparable levels of punishment-induced suppression of cocaine self-administration (suppression ratio: *t*_64_=1.65, *p*=0.10). As previously reported, female rats exhibited more cocaine-induced locomotion than males (Sex; *F*_(1, 31)_=4.91, *p*=0.034, [77]), though CNO (relative to vehicle) in VP-hM4Di did not differentially impact locomotion between sexes (Sex x Treatment interaction; *F*_(1, 31)_=0.33, *p*=0.57) or reinstatement under any condition (*F*<2.5, *p*>0.13). Despite enhanced cocaine-induced locomotion, no sex differences were detected for cocaine-primed reinstatement (Sex: *F*_(1, 22)_=2.06, *p*=0.17), suggesting that cocaine promotes non-specific locomotor activation in females, without enhancing the incentive motivational properties of cocaine under these conditions.

## Discussion

These findings point to a crucial role for VP in cocaine reinstatement following voluntary abstinence, a translationally-relevant model of humans who quit drugs due to mounting negative life outcomes. Indeed, we found that VP plays an especially important role in the most compulsive cocaine-seeking individuals, i.e., the ~20% of rats that tolerated significant footshock punishment to continue taking cocaine. We also found robust reinstatement-related activity in anatomically-defined VP subregions. Our results suggest that unlike connected limbic nuclei, VP plays a critical role in reinstatement regardless of how abstinence was achieved or how relapse was initiated, thereby placing it amongst the most essential nodes within the neural circuits of cocaine addiction.

Persistent drug use despite negative consequences, and long-lasting relapse propensity are cardinal features of addiction in humans [78]. Though compulsive drug intake despite punishment is common in rodents following extended drug access [22, 23, 70], some rats seem to transition to compulsive use even after short access to cocaine. Here, we observed such a subset of compulsive rats, and found that these same animals were also more prone to cue+context reinstatement after voluntary abstinence, similar to prior findings [17, 25]. Importantly, VP inhibition in these compulsive rats had a greater reinstatement-suppressing effect than in punishment sensitive rats, suggesting that VP plays a particularly important role in those rats which pathologically seek drug. VP inhibition only modestly reduced punished cocaine self-administration, but instead selectively reduced hesitations to press the cocaine/shock and inactive levers, which we interpret as reflecting motivation to pursue cocaine, tempered by motivation to avoid being shocked. These results highlight the sensitivity of this novel assay of conflicting motivations during cocaine seeking, and the importance of careful ethological analysis of complex drug-seeking behaviors during such neural circuit manipulation experiments.

Relapse is not a unitary phenomenon, since brain circuits underlying drug reinstatement depend on the drug of choice, mode of abstinence, and relapse trigger [10–12, 26, 79–82]. This said, we show that even under maximally human-relevant conditions VP is broadly implicated in reinstatement regardless of trigger or mode of abstinence. In contrast, other VP-connected limbic regions seem to be engaged differentially during reinstatement after different modes of abstinence. For example, inhibition of basolateral amygdala *decreases* reinstatement after extinction training, whereas the same manipulation during reinstatement following punishment-induced abstinence *increases* drug seeking [10]. These results are consistent with the idea that VP serves as a ‘final common pathway’ of drug seeking [38, 39]. Therefore, VP holds promise as a potential therapeutic target for suppressing relapse in humans, especially since a prior human fMRI report found that activity in the vicinity of VP predicts relapse propensity [83].

Interestingly, in the unpunished safe context, VP inhibition attenuated reinstatement with or without cues, and also after a cocaine priming injection, yet VP inhibition did not reduce cue reinstatement in the punishment context. As expected, conditioned suppression of seeking was observed in this context relative to the safe context, but response-contingent cues nonetheless supported some pressing, reducing the likelihood of a floor effect (Fig. 4). We therefore speculate that VP promotes conditioned drug seeking in a context-gated manner, consistent with prior reports that VP is necessary for context-induced reinstatement of alcohol seeking [44, 45, 49, 84].

VP is heterogeneous, with rostrocaudally- and mediolaterally located subregions, and functionally distinct, genetically-defined neuronal subpopulations [30–32, 54, 60, 84]. We observed broad recruitment of Fos in VP subregions during cue reinstatement tests in both the safe and punishment context, and even in the safe context in the absence of response-contingent cues. This homecage-relative Fos recruitment was most pronounced in rostral, relative to caudal VP. These results suggest a global recruitment of numerous VP subregions during both context- and cue-induced reinstatement, including in the punished context where global VP inhibition failed to suppress cocaine seeking. One possible explanation for this puzzling pattern of effects is that functionally opposed VP cells are engaged in the safe- and punished-contexts, such as the intermingled VP GABA and glutamate neurons which drive appetitive and aversive behavior, respectively [30–32, 61, 62]. Our pan-neuronal chemogenetic approach primarily targeted reward-related VP GABA neurons (~85%), consistent with observed reinstatement suppression in the safe context, where aversion-related glutamate neurons would be less relevant. We speculate that in the punishment context, glutamate and GABA neurons were recruited (explaining Fos results), but inhibiting both concurrently with DREADDs suppressed motivation as well as aversion, resulting in a null effect. More work is needed to parse the specific behavioral roles for VP subregions and neuronal subpopulations in addiction-related behaviors.

The present report firmly establishes VP as an essential node in the neural circuits of translationally-relevant cocaine reinstatement behavior, especially in the most compulsive, addicted-like animals. By better understanding how addiction-relevant behaviors map onto defined neural circuits in the addicted brain, we may reveal neural signatures that could facilitate diagnosis and treatment of addiction in a personalized manner. These and other results suggest that VP plays a key role across relapse triggers and modes of abstinence, making it a promising target for future interventions to treat addiction.

## Acknowledgements

We thank Yavin Shaham for providing MED-PC code for punishment-induced abstinence protocols. We thank Iohanna Pagnoncelli and Stephanie Lenogue for their assistance with behavioral testing.

## Supplemental Figures

**Supplemental Figure 1. Self-administration in male and female rats.** Males (black) did not differ from females (gray) across 14d of self-administration on active lever presses (top), infusions obtained (middle), inactive lever presses (bottom).

**Supplemental Figure 2. Viral expression in hM4Di misses and eGFP controls.** hM4Di misses (thin solid border) and eGFP controls (thick dotted border) depicted on rostrocaudal VP sections.

**Supplemental Figure 3. VP inhibition decreases cocaine-induced rearing, but not distance traveled. A)** Within subjects tests showed that CNO in VP-hM4Di rats fails to alter cocaine-induced horizontal locomotion. **B)** CNO decreases rearing in VP-hM4Di rats. No effect of CNO in control rats for horizontal locomotion or rearing. Control = hM4Di misses and eGFP-only rats. \*\*\**p*<0.001

**Supplemental Table 1.** List of antibodies used for immunofluorescence (columns 1-3) and for immunohistochemistry (column 4).

|  | mCherry+Substance P | GFP+Substance P | Neurotensin+Substance P | Fos+Substance P |
| --- | --- | --- | --- | --- |
| Primary Antibodies | **Substance P**<br>Immunostar (20064)<br>Rabbit anti-substance P (1:5000)<br>**mCherry**<br>Clontech (632543)<br>Mouse anti-mCherry (1:2000) | **Substance P**<br>Immunostar (20064) rabbit<br>anti-substance P (1:5000)<br>**GFP**<br>Abcam (AB13970)<br>chicken anti-GFP (1:2000) | **Substance P**<br>Abcam (ab14184) mouse<br>anti-substance P (1:7500)<br>**Neurotensin**<br>Immunostar (24427) rabbit<br>anti-neurotensin (1:5000) | **Substance P**<br>Abcam (ab14184) mouse<br>anti-substance P (1:5000)<br>**Fos**<br>Millipore (abe457) polyclonal<br>rabbit anti-c-Fos (1:5,000) |
| Secondary Antibodies | **Substance P**<br>Invitrogen (A21206) Donkey<br>anti-rabbit AlexaFluor488 (1:500)<br>**mCherry**<br>Invitrogen (A21203) Donkey<br>anti-mouse AlexaFluor594 (1:500) | **Substance P**<br>Invitrogen (A-21207) donkey<br>anti-rabbit AlexaFluor594 (1:500)<br>**GFP**<br>Jackson ImmunoResearch (703-545-155)<br>donkey anti-chicken AlexaFluor488 (1:500) | **Substance P**<br>Invitrogen (A21202) donkey<br>anti-mouse AlexFluor 488 (1:500)<br>**Neurotensin**<br>Invitrogen (A-21207) donkey<br>anti-rabbit AlexaFluor594 (1:500) | **Substance P**<br>Jackson ImmunoResearch (711-065-152)<br>biotinylated goat anti-rabbit IgG (1:500)<br>**Fos**<br>Jackson ImmunoResearch (715-065-150)<br>biotinylated donkey anti-mouse (1:500) |

**Supplemental Figure 1.**
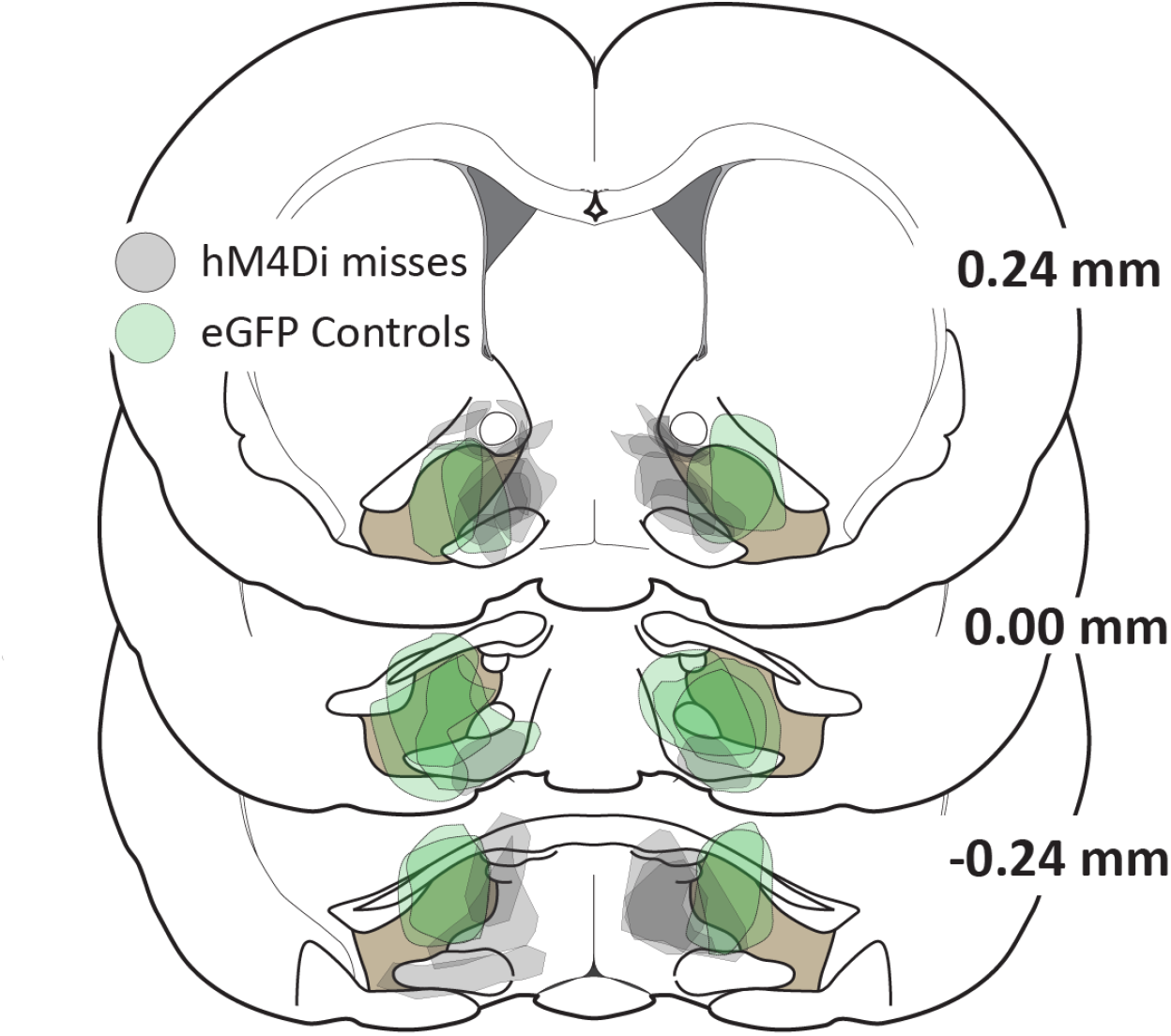
Viral expression in hM4Di misses and eGFP controls. hM4Di misses (thin solid border) and eGFP controls (thick dotted border) depicted on rostrocaudal VP sections.

**Supplemental Figure 2.**
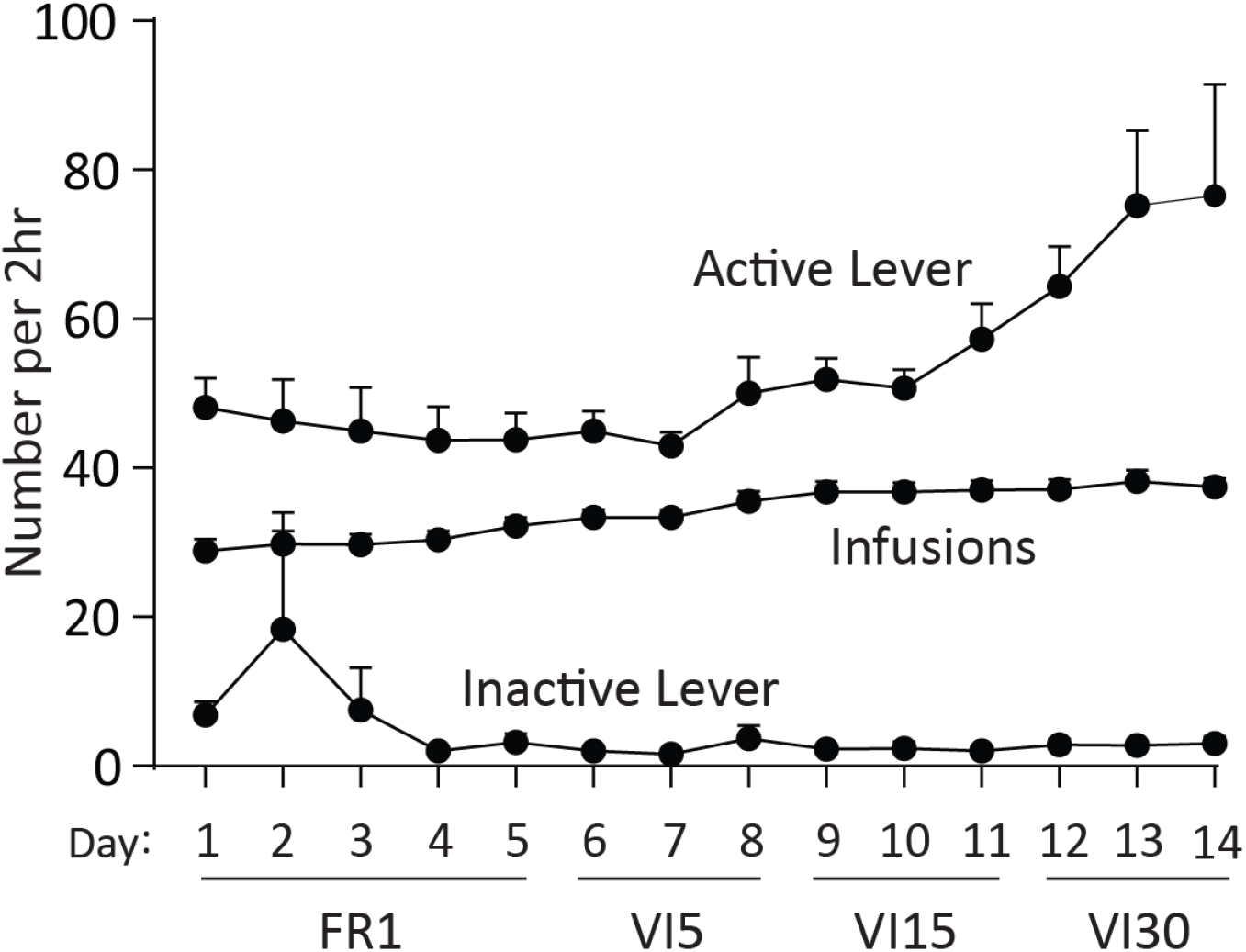
Self-administration in male and female rats. Males (black) did not differ from females (gray) across 14d of self-administration on active lever presses (top), infusions obtained (middle), inactive lever presses (bottom).

**Supplemental Figure 3.**
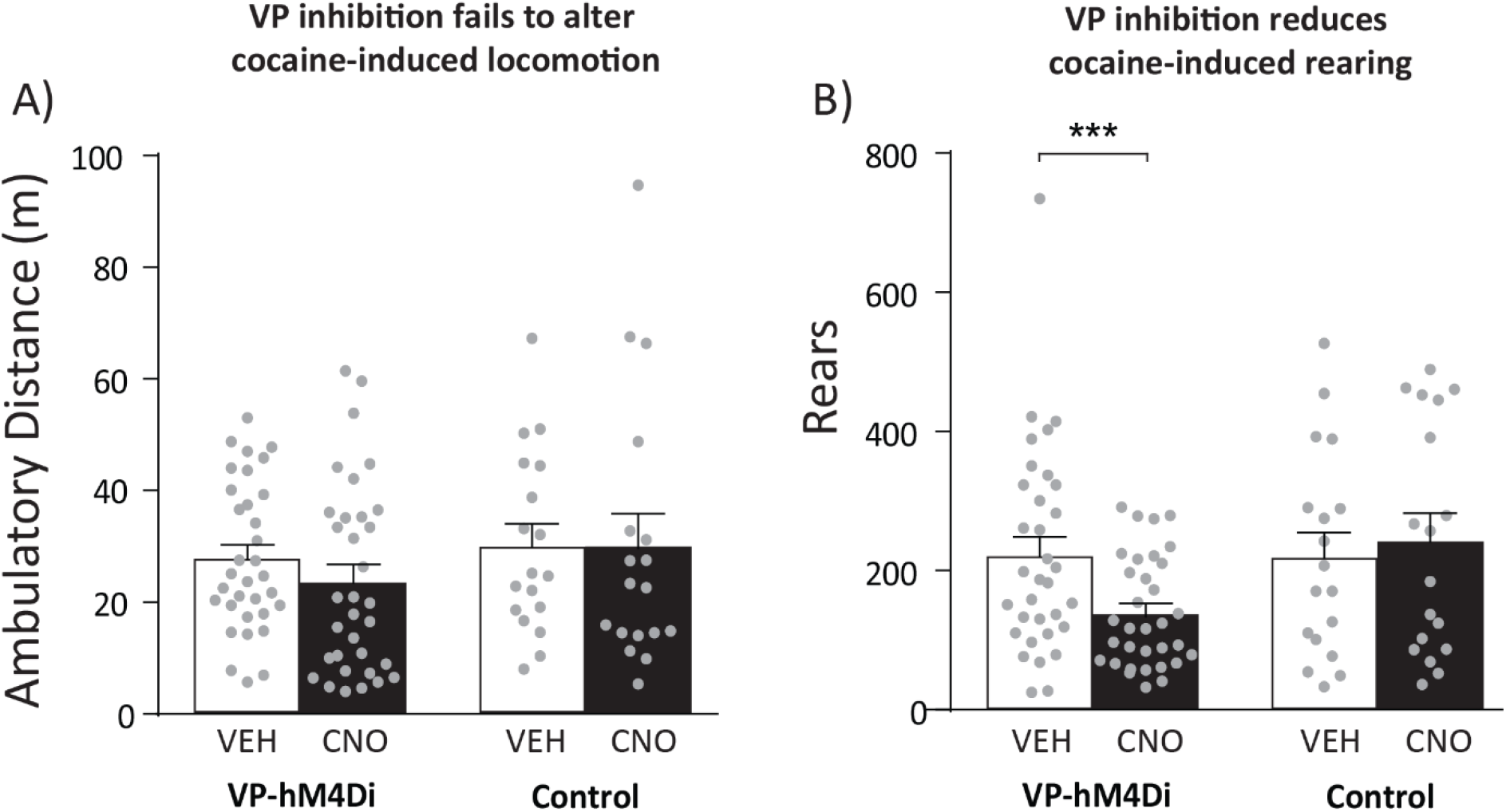
VP inhibition decreases cocaine-induced rearing, but not distance traveled. **A)** Within subjects tests showed that CNO in VP-hM4Di rats fails to alter cocaine-induced horizontal locomotion. **B)** CNO decreases rearing in VP-hM4Di rats. No effect of CNO in control rats for horizontal locomotion or rearing. Control = hM4Di misses and eGFP-only rats. \*\*\**p*<0.001

